## Supplementary Information for "A SUMO E3 ligase promotes long non-coding RNA transcription to regulate small RNA-directed DNA elimination"

### Oligo DNAs

EMA2-C-FW: CCGCTCTAGATTTAATCTTTCTTATGTTAATAATCGTC
EMA2-HA-C-RV: GTCGACACTAGTGGATCCAATGCAAATATTCAAAGAGTT
EMA2-F-FW: GGATCCACTAGTGTCGACTGTTTGTCTTTATCTCGTAGTC
EMA2-F-RW: GGCGCTCGAGTTCTAGGATAACTAATAGAGTGAAC
SPT6-3F-FW2-overlap :TGCTAGCGGATCCGTCGACCTCGAGTCTCTTAGTGTTTTATATAAAACAGC
SPT6-3F-RV1-KpnI: ATTGGGTACCGGGCCCGTATCTTTAACTATCGACAAAACC
SPT6-FW1-SacII: TCCACCGCGGACTTATCACTTAAGGAAGAATATGGC
SPT6-RV2-overlap: ACTCGAGGTCGACGGATCCGCTAGCATGCTCGTTGTCATATGAACGAG
SMT3_HA_FW:CCTTATGATGTTCCTGATTATGCTGGTGGTTCTGGTACTGATTAAAACGC
SMT3_HISEXT_RV: GCTGACCGATTCAGTTCGCTCAATCAGAAAGAGCCACCAACTTGTTC
BTU1_LCFW: AATTTTAATGCGTGTATTTATTTGGGTG
MTT1_5UTR_SeqRV: AGTACTCTATTATGTTTTTTCTTATACAG
EMA2-pBNMB1-FW: ATGATGTTCCTGATTATGCTGGATCCATGAGTTCACTATTTGGAAATAAGAATAA
EMA2-pBNMB1-
RV:TTAGCTGACCGATTCAGTTCGCTCAACTAGTTCAAATGCAAATATTCAAAGAGTTAAGAG
SPT6_FW_Turbo: GATCTGGTGGTTCAGGAAGCGGATCCATGAGTAAACCAAGAAAAAACGAAG
SPT6_RV_Turbo: GCTGACCGATTCAGTTCGCTCAATCAATGCTCGTTGTCATATGAACGAG
SPT6_N_FW_HA: GTATCCTTATGATGTTCCTGATTATGCTGGATCCATGAGTAAACCAAGAAAAAACGAAG
SPT6_N_RV: ATACTTTGCGCCATAATCACCTAC
SPT6_5F_FW: CAAATTTTTACTGGAAAAATGCAGCTAAATTATGAGACCAAACTAAG
SPT6_5F_RV_HA: GCATAATCAGGAACATCATAAGGATACATTTTTCAAATATTTGCTGTGAAGACT
SPT6-KO-5F-FW: CGAATTGGGTACCGGGCCCCCCCTCGAGCATAATATCATTCTCCACTCAC
SPT6-KO-5F-RV:
CAGTAAAAATTTGGATCCTATCGAATTCCTGCAGCCCTGCTTAAAAGTGCCTATGTTAGC
SPT6-KO-3F-FW: CGATAGGATCCAAATTTTTACTGGAAAAATGCAGCCCTCTCGTTCATATGACAACGAG
SPT6-KO-3F-RV: GGCGGCCGCTCTAGAACTAGTGTATCTTTAACTATCGACAAAACC
SPT6_M249_FW: CTTATGATGTTCCTGATTATGCTGGATCCATGTAGGGACGTAAAGCTCCTGATCC
SPT6_M249_RV: GTTCGCTCAATCAATGCTCGTTGTCATATGAACGAGATCTAGAGCG
M5'-3: AGGTACGATAGATCGACTGACGG
M5'-4: AATAATAAGGAACCTCTTACTGTG
M3'-3: ACTAAAATATTTATCTTCTTTTCTGC
M3'-4: ACTTTCAAAAAATTAATTTTGAGTAAAG
L8_5'-1: ATTTGCTTTTAGGAAGAGATATCAC
L8_5'-2: ACATACATTTGAAATATCACCAAGC
L8_3′-1: ACAAGAAAGGTATTCATTCATTCCTC
L8_3′-2: TGATAAAAAAGTGGTGAAAAAAGGAC
R2_3'-1: TAATTTTAGGGCGAATCACC
R2_3'-2: ACTCATAAATAAAATCATATCATAGTCTAG
R2_5'-1: TTAGCAAAGTGCATTAACCTC
R2_5'-2: TCTCTTGAAATTGGGCAAAAACTG
RPL21-FW: AAGTTGGTTATCAACTGTTGCGTT
RPL21-RV: GGGTCTTTCAAGGACGACGTA
NMC1_f53600: GTGCAATTTTACACCGTCAGG
NMC1_r55732: AGAATCTAACTCATAGCACTGG

### Synthetic DNAs

>EMA2-Ec
CGATCGGGATCCATGTCTAGTTTGTTCGGTAACAAGAACAAGGAAGTTATCTTCAATTTTTGCCAAAAGTCTCA
GGAAGAAGATCAAGACATGTTTGCGGACCATGAAATTCTGATTGATGAACAATTACAGTGTTTTAACTGTAACA
AGTACTTTGATCCCACGAAGTACGGGATCGATTTGTTAGACGAGATGGACTTCAACAAGAATCTGGACACTTTT
TGTTGTGAGAATTGTTATCTTTCGTTGCATTTTCCTTTTTATTATCAATTGGATTTATTGTTGAAAATGTCAAA
CCTGGAAGTGAACACGAAGTACGAATTCAACCTTGTACTGCAGGAGGAGTGCTACCAAGACAGTTGCATCCTGG
GCGTGTTCTTAAGCAAGCAAAATCTGATTAATGTCTTCAAGAAACAGACGGGGAGTCATAACTTCATCTTAAAC
GTGAAAATGAATGGGCAAGACCCCTCCCAGGAAGGCAGTAACTTTTTTGTGATCGACTCGGATGACTTAACGCA

GATGCAAAATACTTTGAACTTCGAATGTCTTAACTTTTCCTTTCAGAAATCTTTCATCTCGAACATCATCTTAA
TGAACGACCAACAGAAAAGTCAGCTTAAAACGGCCTCTCCGAACATCATCAGCGTAATCCTGCACTGCCGCAAA
ATTCGTGTCTCTGCGTATGTCGCGAAACTGTTTAAGAAGAAGTTCTTTAACTGTTTAAATCATGAAAGTTTATT
CAAGGTTTTCGAGGAACAAGAGCAAAAATATAAGCAAAAAGACCAGATCCAGAAAATCAAATTACAAACCAAAC
AGAACTCGCTTTCAACGTCTCAAATTCATCAGTCTAGTACCCTGAACGAAATTCAGCCCAGCCAGCTTCAAACC
CATAAGGAAACTTCGAACATCTCGCAAAGTAGCCAGCTTAACAAAGATTTCTGTGAAAAAAAAGAACTGACCGA
GATCTATCTTCGCATTAAAAACTTTCACCAGATGTTCCCGTTCATGTACAAGAATGAGATGGTGGTTAAGATGG
ACGATAACGCAGTTATCCAATACCCCGCGTTTACCTACCATCACTCGCACCCCTTTGACTTACGTGATTACTGC
TACCTTAACCAGGTGAATCCTACCTGGTTGTGTCCCATCTGCAACAAGCACAAGATTTTCTTGAAGGAGATCCA
GCTGGATTTTTATTTGTTTGCGTTGATTCAAACACAGTATCTTAAGGACTCCTATATTTCGGACAAAATCCAGC
AGATTCTTCAGTCGAAGCCACAGTTTGATTCTGAATTTTTTAGCAATCAACTGAAACAGCTGAACTTGCACTCA
ATGAAGGTACTGGAGCAAACACAGCAACGCACCCAGTCTAACGAGATTAAGATCCAGCAGAACTTATCCAATAA
GTTGAACCAGACTAACAAGAATTTTATCCAGATCTACAAAAAGTCGAATCAGTTGGGGCAAAAAGAGAATATTC
TGAAGAACAGTACTAAAATTAAGCAATTTAAGTGTAAGTCATCTATTGATCAGAATACAAATTATGAACAGCGT
AACCACAAGAATGGGAATCAAAATCAAAAGTCAAAAACCTCAGAGTTTTTCAATGAAAATTTGAACTGTACTGA
CTTAAGCCAAGACAATTACCTGGATTTCATTCAGGATTTTGACGAATTTCAATCTTTCTTAAATCCTTACCAGA
AGCAACATTCGATGCGCAATAGCAAAGATATTTCTGTACGTCATAATATGGATCAAGCTAACCTTATCGACGAG
ATGTCTATCCTTAACATTCAGTGCAATGAAATCGCCTCCCAAAAATCTTCAACCCGTGTTTACAGCAAATTAGT
AAACAAATCTAAATCGAAGACCTCATCTTCTAACAAAGAAGAGAAGCAGAAGCAGAACGACGCATTTAAAGAAA
ACCAAATCGACATTGAGCAATCAGGCCAGATTATCGAAGAAAAAGTCCAAGAGAAACCGAGCTTGAACGATCAA
ATTGATGCTAATAAGAACATCCAGTTCTCCCAACAAATTGAGTCCTTTTCCGACCTGAGTCAGAAGTTCATCTC
CCAGCAAAAAGAAGAGCGTACTCTTGAGGTTGCAAATCAATTCTACTTGGACACTTCGTTCCTTCGCTCCCCTC
TTGAAATCAACGTCTTGTTTGACGTATCCTATTTGCAATTCAATAAAGAAGAGCAGGAGTATCAGTCAGTTCAG
CAGATCATTCAGCTGATTGACTTCCAAGAAGACAAGCAGATGTTAGAGAAGCTTTTCAACATTGACATCCTTCA
AAATCACTATTTTATTTTCAATATTTTCAATCTGAGCTATGTAAATAATCGTCAGTATCAAGACTTGCACACAA
TCTTAAACCAAAACCCTTCGACTAAAACTTTTTTTCAAGTCTTGATCAACGTTCAAGAGCAGTTCATCAACCTT
GGTATGAGTTCAGAGTTTTATACGTTAATTTCTCAGTTCTATTATAAAGTTATGTCTAATAAGCAGCGCAATAA
TAAGATTTCTTACGACCTTTTGGATTCCGAAAGCCAAATTACGTCGCTTAATTATTCGCAAAACAAAAACTTTT
TAGAGAAACTTATTGAATGCTTCCTGGGAATGTCGATTTTTAAGCAACTGTCCAACGAAGATGTCATTGAAAAT
TTCCAACAAATGGTCTTTGAATACTTATTGTTCAAGCAACAAAATATCCCGACTTTGATTAATATTCACAATAT
CATCATCAACTATTTTCCCAATCAATGCCTTAACTTAATCGAATCGATCAAGAAATCTTTGAATAGCCTGAACA
TTTGCATCTGACTCGAGCCCGGG
>UBC9-Ec
GGCCTGGTGCCGCGCGGCAGCCATATGATGCAACAGCAGAATAAAGAAGTTAACGAACTTGTAATTACGCGCCT
GAAACAGGAGCGCAAACAGTGGCGTCAGGATCATCCTCATCGTTTCGTCGCCAAACCCATGACTAAAGAAGACG
GTACCATCAATATGCTTAAATGGTACTGTGAGATTCCGGGACCGGAAGGGTCACCCTGGGAGGGGGGCGTTTAC
ATTTTATACATGGACTTTTCACAAAGCTACCCTATTAAACCCCCTAAGTGCCAATTCAAACCGGTCCTTCCTCA
CCCCAACGTGTATCCAAGTGGCACCGTCTGCTTATCAATCCTGAACGAAGAAGAGGACTGGAAATCTGCAATTA
CGATCAAACAGATTTTGATGGGAATCCAAAAGTTACTGAAGGATGAACCGAACATTGACAGCCCTGCCCAGTAC
GAGCCCTGTGCTTTATATCGTAGTGACAAGGAAAAATACTACAAAAAGGTACGTGAGTTTGCTGAAACTATGAA
GAAGAAGGATtgagGATCCGAATTCGAGCTCCGTCGACA
>SPT6-DOL-KtoR
CCTTATGATGTTCCTGATTATGCTGGATCCATGTCTAGACCTCGTCGTAACGAGGAGGTTGATGATCAGATGAG
AAACATCGGAGATGAAGAAGAAGAAGATTATCAAGGACAGGTCGAGGAAGAAGAAGATGGAAACAATCAAGTCG
AGGAAGACGACGAGGAAGAAGAGGATGATGACGAACAAGACGATTACCAACAGGACGGATTTGTCGTCGGAGAT
AGCGAAGAAGAGGAGATAGAAGAGGAAGAAGATGAAGATGCCGAAGAGAGACGTCGTCGTAGAAGAGAACGTAG
AAGACAAAGAAGATTGGAGAGACAAAACCAGCACAGAAGAATACTCAGACGTGGTCCAAGACGTCGTGAGTTGA
GCGAGGATGAAATGGAGAACAATGAGATTCAAGATTTAGAAGAGTATCATGAAGACTCTGAAGAGGGTATTCCT
GACCGTCGTAGAAACCAGCAGTCACGTACAGGAAACGACGGTCACCAGGGAGACAGAATGGACCTTGAGGACCA
AGGTAATGAGGACTATCTTGAGGACGTTGGTGACTATGGAGCTAGATACGGAAGAACAGCCTTCGAAGAACTCT
TTATGGATGATGAGTCAGGTGAAGAGGAACAGAACGAGAATGAAGACGACGAGAACCGTGAAGTAGACGACTAC
ATCGACGTCCAACAGTTGTTCGAACCCGATGACTTGCGTAGACGTTTCGAAAGAGACGACGACCGTAGAATTCG
TGAAGAGGATATACCCGAGCGTCTCCAAATCCGCATGTAGGGACGTAAAGCTCCTGATCC
>SPT6-118KR-gBlock-1
CCTGATAGAAAGAGAAATCAGTAAAGTAGAACCGGTAATGATGGTCATTAAGGAGACAAGATGGATTTAGAAGA
TTAAGGTAATGAAGATTACTTAGAAGATGTAGGTGATTATGGCGCAAAGTATGGAAAAACAGCCTTTGAAGAAC
TTTTCATGGACGATGAGTCAGGAGAAGAAGAGCAAAATGAAAACGAAGATGATGAAAATAGAGAAGTTGACGAT
TACATTGATGTGTAATAGCTTTTTGAACCTGATGATCTTAGAAAGCGTTTCGAGCGTGATGATGATAAAAAAAT
TAGAGAAGAGGATATACCTGAGCGCTTATAGATTCGTATGTAGGGACGTAGAGCTCCTGATCCAGAAGAGCTCA
TGGAAGAGACAAGATGGATATCAGAAAAAATTAGGACTATAAGGGATTATAGGGGTAGCAGGAGGTTGAATGAT
ATTAATTTCCATTCAAAAATATTCTCTTTCCTCCAATCTCTACATTTGGATAAAGAAGAAGTTATGTATATCTA
TACATACAAGAGGAATGAATTTCATCCACAGTTTGATCTTGAAGACTTATGGAGATTATATGACTTAGATGGCG
AATGGGCTCAATTCAATAGACAAAAAAATCGCATTCTATTATAAATTTAATAGCTTAGAAGAGAACTTGAGAAT

ACTCCTCTGCTAATTTAAAATAATATAATTGATTTTGAATAAGTTAGAAGAGTTGAAGAATTTTTCCACAGAGC
CATTGATACTTAATCTCTTAAGTTTATCAAGGAATACTTTGATTACCTATTTATCAAAATTTATCCTTAAAGGG
AGCACTAAAGATTAAGAATTCGTAGAGACCGTAAAACTAGAATGATTAACACTTATATTTAATGTAGGGTTCAT
AGGTTAGTTAGTACTTTAACTCTTTCTCCTTAGGATCTCGTTAGAAACTTAGAAGAAATGAACTAAATTAATTA
ACCTCAATTACAACAAAATAGGCCCTAAGCCTATGCTGATAATTTAATTTCAGAAAACCCAAATGTTCCTTGTA
TGAGAGAAGCTATAGAGGCTTTAGAATGCATGGTTGATTATCTATCAAATGAATTATTTAACTATCCTCCCCTT
AAAAGATGGTATTATGAAATGTTTAGATAAAGATGCTTAATTTCTACTGAACCCACAGATCTTGGTAGAAGGTA
AATAGATATGTTCCATCCCCTATACCCAGCTAAAAGAATTCATAGACGTCCTTACACTTCTTTCACTGATGACA
CTTGGTTGCTCTTAGAAGAAGCTGAAAGATTGAACTTTATTACAGTTAGGATTTTCTTAAGAAGTGAAAATGAT
AATAACTCAGACAATAACAGCATATTTGATATTATGAGTAGAAATTACATGATCAGTTTAAGAAATTTAGAAGC
TCAAATTAGACAAATTAATTCTGAATGGAATTTCTTCCGTAGATACAACATAGAAAATGTTTGTGATAGGTTTT
TCTTACCT
>SPT6-118KR-gBlock-2
CATAGAAAATGTTTGTGATAGGTTTTTCTTACCTTATGCCTAGAGCTTAGTAAGAATAGAACTTCATGAAAATG
CTGAAAAGTGGGTTGTATCTTAATGCTAACAAAGGTTTTACAACATGATTAATGTTGAACCTTAGAGAAATGGT
AGAATAATGTCAGTTGTCCAAGACACAAATGGTAGAATAGGTGTTGTCTTTGTTGATGAAAATGGAATTCCTAA
TAATTCTATGATTTTGAACTACTTCACCAGGAGAGAAGACTAGCTTTCAACTAGAGCCAGACTAGAAAAAACCT
AAGAAGAACATGAACTTGAAATTAGGTTTAGATCTTTTGGACCTTAATTAATTGTTGTTGCTGCCAATAGCATA
GAATCTTAAAGACTTAGAAATCTACTTCATTAAAAATATGAAAGATCTTAATCTTATATTTAATATGGAGATGA
CACAATCCCATAAATTGTGGCTAAGTAACCTACAATTCAAGGACCAGGAAATACCTTACAATTTAGAGATTCTG
TACTCAGAGTTGCTCTTTCATAAGGAAGAATACTTTTAAATCCTACTGCAGAAATACTCTCGCTTTGGAACGAT
AATGTTTAACAAAACGGGTGTCTTCATATCCCCCTTCATCCATCTTAAGAATATATTAGTCTTGATAAGCTCTA
GTACTCACTAGAGAAATGTTGTGTTTAAATAGTCAATTTTATGGGTGTTGAAATGGATATGCTTTAAGATTAGC
CTCACCTTAGACACCTTCTATAACACGTATGTGGTTTAGGACCTAGAAAGGCATTAAGGCTAATTGAGCTCTTA
GAAAGAAATGCTGTAATTTTTGAAAGAGAAAGGGAAGTACCTAAAAGGAGAGGTCTATTAATTACTCAACTTGA
TATAAGGGATGTTGTATTCAGGAATATTAATGGTTTCATCAGATTTAGATTTTCAAAAGACCCTATTGAAAAGA
CTAGAATCAACAGAGATTTATACAGAATTGCAAGAAGGATTTGCAGAGACAGTGCTGATAACTTCTAAATTAGA
AGCTAAAGTGATGAAGATATAGTAGAATATGTTATGAGAAATCCCAGATATATTGATGCTATGGACTTGGAAGA
TTATGCTTAACAGCTTGAGGAGATGAGAAATCAACCAAACATGGCTCCTGTATTAGATTTTGTCAGAGATGAGC
TTATCAACCCATTTAGCTATAGAAGAAATACCTATGAAGCAATTAGCGATTAAGACTTATTCTTCAAAATGATT
AGAGAAAGTCCTCAAACATTCCGTAAAGGAATGATTGTTTCAGCAAGAATCATTCAAATTAGACCTAGAGAAAA
TCAATAAATAAGACAACACTAACTACTTGTTAGGATTGTTGATAACGACCTAAGATCTAGCATACTTGTTAGTG
AAGAGGAATCAAGGAATTATAGGGTTGGTGATATTATTAAAGCTTATGTTGATCAGATTTTTGTCCTTGACAAC
AGGCGC
>SPT6-118KR-gBlock-3
GATTTTTGTCCTTGACAACAGGCGCTCCAAGGATAGAATTATTGGTTTCGATGTCAATTGTGTGACTTTAACTT
TTAGAGCCATCAGATTTAATGATTTAGTCAGAGAAATGAGATAATTTTAAGACTTAGATATTCTTAGTACTTTC
AGGTTCATTGAAGCTGAAGACAGGCCATTGGGTATTGATGTCACTGAAAGAAGAGCTAGGAGATTCGAACCAAG
AAGGATTGCTCATCCTAATTTTAGAAATATAAGTATTACAGATGCTGTAAGGCTTTTGAGGAATGCAAGAAATG
GTGAATTTATTATTAGACCCAGTTCAAAAGGTAGACAATACCTTGCCATAACATGGAGATTCTTCGATGACGTG
TTTGTCCACTTATCACTTAGGGAAGAATATGGCAGAGAAAGGGGGTTCTAAACGAGATATGTTTTAAATGACAG
AGAATCGTTTGATAACTTTGATGAAATTATTGAAAGATATATCATTCCTTGCAATAATCACATGAACTCAGCTA
GAGATAATCGTAGGTTTTCAAGAAGATCAATAGAAGAAATAGAATAGGAGCTAAGGAGAAATAGAGAAGAATAA
CCAGACATTATTCACTATAACTTTTGTTGTGTCCCTAGATATCCTCAATTTATCGTCTTGCTATATTGCTCTAG
ATAGGATTAAGTCACCAAAGAGTGGATTAAAGTTAAGTATTAGGGCTTTTATTTCCATGAAAGGTATTTTGGAC
AATTAAGGGATTTAATTAGATGGTTTAGAGATTGCTTCCATACTCCCGAATACAGAAGGTATGTAAAAAGAGCT
GAAGAACCATATGCATCTAGGACCCCTAGCTCATTTAGTAATTAAGGAACTATTGGCATAAGATAAGAAGGTGG
TGTCAGATATGAAAGATCAGAGAGAGGCATTGGAGGACAAAGATGTTAAAACTGTGGTAGATATGGTCATGCTG
CTAATAACTGCAGAAATAGAAGATCCTATAATTCCTCTTCAGGATAGGGCAGTGCATCTGGAGCAAAACAATGT
TTCCACTGTAGGGGTACTGACCACTTTATTAGAGACTGCCCAAATAGAACTCATCAAAGCAATAGAGGAAGTGG
CGGTTTTAGCTATAGCAGAAGAGATAGAAATCAATAACACTCACGCTCTAGATCTCGTTCATATGACAACGAGC
ATTGATTGAGCGAAC
>SPT6-42KR-gBlock-1
CCTGATAGAAAGAGAAATCAGTAAAGTAGAACCGGTAATGATGGTCATTAAGGTGATAAGATGGATTTAGAAGA
TTAAGGTAATGAGGATTACTTAGAGGATGTAGGTGATTATGGCGCAAAGTATGGAAAAACAGCCTTTGAGGAGC
TTTTCATGGACGATGAGTCAGGAGAGGAGGAGCAAAATGAAAACGAAGATGATGAGAATAGAGAGGTTGATGAT
TACATTGATGTGTAATAGCTTTTTGAGCCTGATGATCTTAGAAAGCGTTTCGAGCGTGATGATGATAAAAAAAT
TAGAGAAGAGGATATACCTGAGCGCTTATAGATTCGTATGTAGGGACGTAAAGCTCCTGATCCAGAGGAGCTCA
TGGAGGAGACAAGATGGATATCAGAGAAAATTAGGACTATAAGGGATTATAGGGGTAGCAGGAAGTTGAATGAT
ATTAATTTCCATTCAAAAATATTCTCTTTCCTCCAATCTCTACATTTGGATAAAGAAGAGGTTATGTATATCTA
TACATACAAGAGGAATGAGTTTCATCCACAATTTGATCTTGAGGACTTATGGAGATTATATGACTTAGATGGCG
AATGGGCTCAATTCAATAGACAAAAAAATCGCATTCTATTATAAATTTAATAGCTTAGAAGAGAGCTTGAGAAT
ACTCCTCTGCTAATTTAAAACAACATAATTGATTTTGAGTAAGTTAGAAAAGTTGAGGAGTTTTTCCACAAAGC

CATTGATACTTAATCTCTTAAGTTTATCAAGGAGTACTTTGATTACCTATTTATCAAAATTTATCCTTAAAAGG
AGCACTAAAGATTAAAAATTCGTAAAGACCGTAAAACTAGAATGATTAACACTTATATTTAATGTAAGGTTCAT
AAGTTAGTTAGTACTTTAACTCTTTCTCCTTAGGATCTCGTTAGAAACTTAGAAGAGATGAACTAAATTAATTA
ACCTCAATTACAACAAAATAAGCCCTAAGCCTATGCTGATAATTTAATTTCAGAAAACCCAAATGTTCCTTGTA
TGAAAGAAGCTATAGAGGCTTTAGAATGCATGGTTGATTATCTATCAAATGAATTATTTAACTATCCTCCCCTT
AAAAAATGGTATTATGAAATGTTTAAATAAAGATGCTTAATTTCTACTGAACCCACAGATCTTGGTAAAAAGTA
AATAGATATGTTCCATCCCCTATACCCAGCTAAAAGAATTCATAAACGTCCTTACACTTCTTTCACTGATGACA
CTTGGTTGCTCTTAGAAGAAGCTGAAAGATTGAACTTTATTACAGTTAGGATTTTCTTAAGAAGTGAAAATGAT
AATAACTCAGACAATAACAGCATATTTGATATTATGAGTAAAAATTACATGATCAGTTTAAAGAATTTAGAAGC
TCAAATTAGACAAATTAACTCTGAATGGAACTTCTTCCGTAAGTACAACATAGAAAATGTTTGTGATAAGTTTT
TCTTACCT
>SPT6-42KR-gBlock-2
CATAGAAAATGTTTGTGATAAGTTTTTCTTACCTTATGCCTAGAGCTTAGTAAGAATAGAACTTCATGAAAATG
CTGAAAAGTGGGTTGTATCTTAATGCTAACAAAAGTTTTACAACATGATTAATGTTGAACCTTAGAAAAATGGT
AAAATAATGTCAGTTGTCCAAGACACAAATGGTAGAATAGGTGTTGTCTTTGTTGATGAAAATGGAATTCCTAA
TAATTCTATGATTTTGAACTACTTCACCAGGAGAGAAGACTAGCTTTCAACTAAAGCCAGACTAGAAAAAACCT
AAGAAGAACATGAACTTGAAATTAGGTTTAAATCTTTTGGACCTTAATTAATTGTTGTTGCTGCCAATAGCATA
GAATCTTAAAGACTTAGAAATCTACTTCATTAAAAATATGAAAAATCTTAATCTTATATTTAATATGGAGATGA
CACAATCCCATAAATTGTGGCTAAGTAACCTACAATTCAAGGACCAGGAAATACCTTACAATTTAAAGATTCTG
TACTCAGAGTTGCTCTTTCATAAGGAAGAATACTTTTAAATCCTACTGCAGAAATACTCTCGCTTTGGAACGAT
AATGTTTAACAAAACGGGTGTCTTCATATCCCCCTTCATCCATCTTAAGAATATATTAGTCTTGATAAGCTCTA
GTACTCACTAGAGAAATGTTGTGTTTAAATAGTCAATTTTATGGGTGTTGAAATGGATATGCTTTAAGATTAGC
CTCACCTTAGACACCTTCTATAACACGTATGTGGTTTAGGACCTAGAAAGGCATTAAGGCTAATTGAGCTCTTA
GAAAAAAATGCTGTAATTTTTGAAAGAGAAAAGGAAGTACCTAAAAGGAGAGGTCTATTAATTACTCAACTTGA
TATAAAGGATGTTGTATTCAAGAATATTAATGGTTTCATCAGATTTAGATTTTCAAAAGACCCTATTGAAAAGA
CTAGAATCAACAAAGATTTATACAAAATTGCAAGAAAGATTTGCAGAGACAGTGCTGATAACTTCTAAATTAGA
AGCTAAAGTGATGAAGATATAGTAGAATATGTTATGAAAAATCCCAAATATATTGATGCTATGGACTTGGAAGA
TTATGCTTAACAGCTTGAGGAGATGAAAAATCAACCAAACATGGCTCCTGTATTAGATTTTGTCAGAGATGAGC
TTATCAACCCATTTAGCTATAAAAGAAATACCTATGAAGCAATTAGCGATTAAGACTTATTCTTCAAAATGATT
AGAGAAAGTCCTCAAACATTCCGTAAAGGAATGATTGTTTCAGCAAAAATCATTCAAATTAGACCTAGAGAAAA
TCAATAAATAAGACAACACTAACTACTTGTTAGGATTGTTGATAACGACCTAAGATCTAGCATACTTGTTAGTG
AAGAGGAATCAAAGAATTATAAGGTTGGTGATATTATTAAAGCTTATGTTGATCAAATTTTTGTCCTTGACAAC
AAGCGC
>SPT6-42KR-gBlock-3
GATTTTTGTCCTTGACAACAAGCGCTCCAAGGATAAAATTATTGGTTTCGATGTCAATTGTGTGACTTTAACTT
TTAAAGCCATCAAATTTAATGATTTAGTCAGAGAAATGAGATAATTTTAAGACTTAGATATTCTTAGTACTTTC
AAGTTCATTGAAGCTGAAGACAGGCCATTGGGTATTGATGTCACTGAAAGAAGAGCTAGGAGATTCGAACCAAG
AAAGATTGCTCATCCTAATTTTAAAAATATAAGTATTACAGATGCTGTAAGGCTTTTGAAGAATGCAAAAAATG
GTGAATTTATTATTAGACCCAGTTCAAAAGGTAAACAATACCTTGCCATAACATGGAAATTCTTCGATGACGTG
TTTGTCCACTTATCACTTAGGGAAGAATATGGCAAAGAAAAGGGGTTCTAAACGAAATATGTTTTAAATGACAA
AGAATCGTTTGATAACTTTGATGAAATTATTGAAAGATATATCATTCCTTGCAATAATCACATGAACTCAGCTA
AAGATAATCGTAAGTTTTCAAAAAAATCAATAGAAGAAATAGAATAGGAGCTAAGGAGAAATAGAGAAGAATAA
CCAGACATTATTCACTATAACTTTTGTTGTGTCCCTAAATATCCTCAATTTATCGTCTTGCTATATTGCTCTAA
ATAGGATTAAGTCACCAAAGAGTGGATTAAAGTTAAGTATTAGGGCTTTTATTTCCATGAAAAGTATTTTGGAC
AATTAAAGGATTTAATTAAATGGTTTAAAGATTGCTTCCATACTCCCGAATACAAAAAGTATGTAAAAAGAGCT
GAAGAACCATATGCATCTAGGACCCCTAGCTCATTTAGTAATTAAGGAACTATTGGCATAAGATAAGAAGGTGG
TGTCAGATATGAAAGATCAGAGAGAGGCATTGGAGGACAAAAATGTTAAAACTGTGGTAAATATGGTCATGCTG
CTAATAACTGCAAAAATAAAAGATCCTATAATTCCTCTTCAGGATAGGGCAGTGCATCTGGAGCAAAACAATGT
TTCCACTGTAAGGGTACTGACCACTTTATTAAAGACTGCCCAAATAAAACTCATCAAAGCAATAAAGGAAGTGG
CGGTTTTAGCTATAGCAGAAGAGATAGAAATCAATAACACTCACGCTCTAGATCTCGTTCATATGACAACGAGC
ATTGATTGAGCGAAC
